# Integrated Proteomic and Metabolomic Analyses of the Mitochondrial Neurodegenerative Disease MELAS

**DOI:** 10.1101/2021.12.18.473301

**Authors:** Haorong Li, Martine Uittenbogaard, Ryan Navarro, Mustafa Ahmed, Andrea Gropman, Anne Chiaramello, Ling Hao

## Abstract

MELAS (mitochondrial encephalomyopathy, lactic acidosis, stroke-like episodes) is a progressive neurodegenerative disease caused by pathogenic mitochondrial DNA variants. The pathogenic mechanism of MELAS remains enigmatic due to the exceptional clinical heterogeneity and the obscure genotype-phenotype correlation among MELAS patients. To gain insights into the pathogenic signature of MELAS, we designed a comprehensive strategy integrating proteomics and metabolomics in patient-derived dermal fibroblasts harboring the ultra-rare MELAS pathogenic variant m.14453G>A, specifically affecting the mitochondrial respiratory Complex I. Global proteomics was achieved by data-dependent acquisition (DDA) and verified by data-independent acquisition (DIA) using both Spectronaut and the recently launched MaxDIA platforms. Comprehensive metabolite coverage was achieved for both polar and nonpolar metabolites in both reverse phase and HILIC LC-MS/MS analyses. Our proof-of-principle MELAS study with multi-omics integration revealed OXPHOS dysregulation with a predominant deficiency of Complex I subunits, as well as alterations in key bioenergetic pathways, glycolysis, tricarboxylic acid cycle, and fatty acid β-oxidation. The most clinically relevant discovery is the downregulation of the arginine biosynthesis pathway, likely due to blocked argininosuccinate synthase, which is congruent with the MELAS cardinal symptom of stroke-like episodes and its current treatment by arginine infusion. In conclusion, we demonstrated an integrated proteomic and metabolomic strategy for patient-derived fibroblasts, which has great clinical potential to discover therapeutic targets and design personalized interventions after validation with a larger patient cohort in the future.

Graphic Abstract:
Integrated proteomics and metabolomics of patient fibroblasts revealed dysregulations in arginine biosynthesis, OXPHOS complexes, and bioenergetic pathways in MELAS, a mitochondrial neurodegenerative disease caused by mitochondrial DNA mutations.

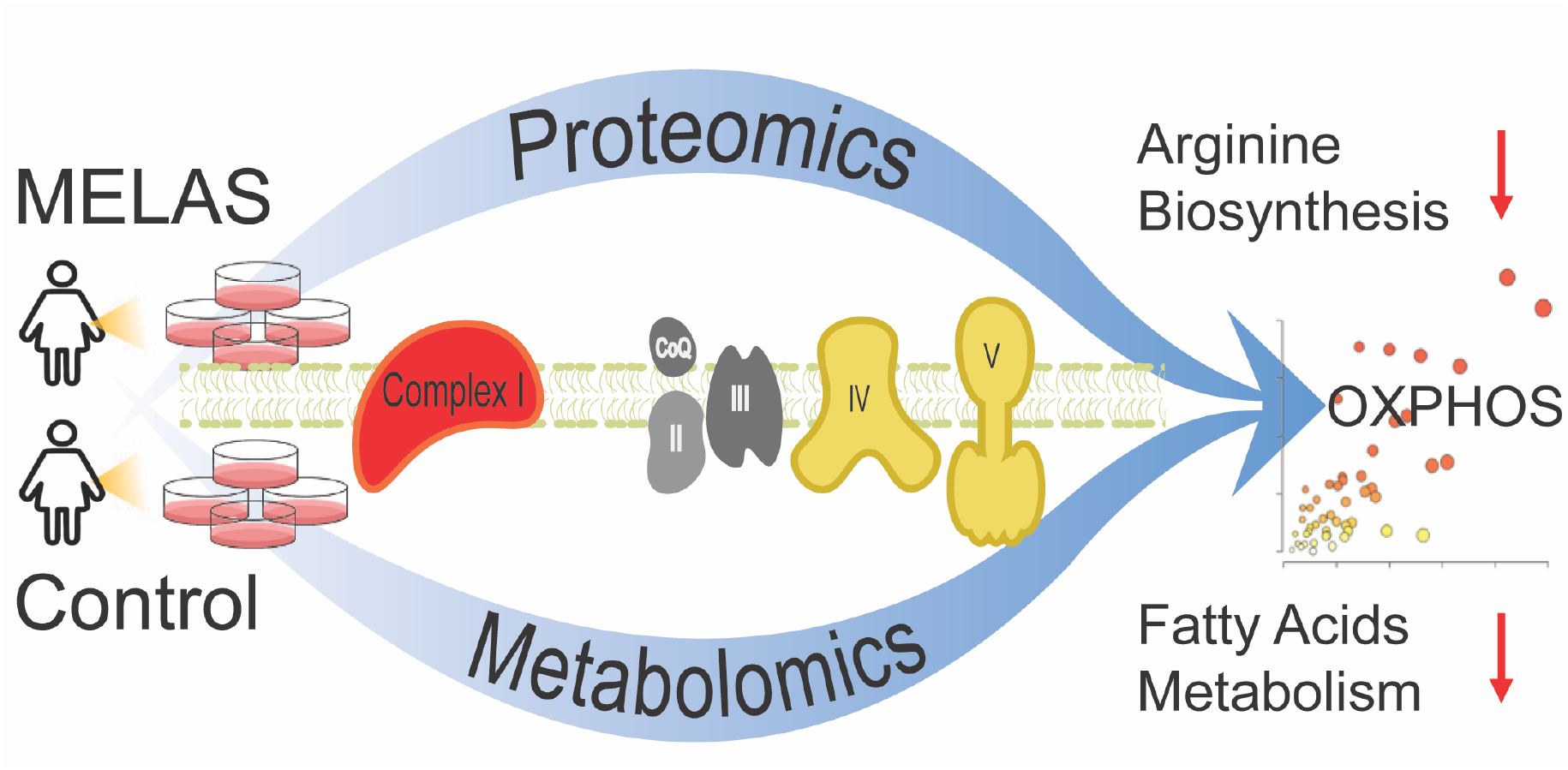

## INTRODUCTION

The maternally inherited mitochondrial disease MELAS (mitochondrial encephalomyopathy, lactic acidosis, stroke-like episodes syndrome) is a progressive neurodegenerative disease with great genetic and clinical heterogeneity ^1^. MELAS has a predominant childhood onset with no gender bias. The neurological features of MELAS include stroke-like episodes, encephalopathy with seizures, lactic acidosis, hearing loss, myopathy, neuropathy, tremor, cognitive defects, and dementia. Non-neurological symptoms also present in MELAS patients, such as cardiomyopathy, nephropathy, diabetes mellitus and gastrointestinal ^1^. MELAS is caused by mitochondrial pathogenic variants affecting the oxidative phosphorylation (OXPHOS) pathway, responsible for ATP synthesis, thereby leading to a chronic energy deficit. About 80% of MELAS patients harbor the maternally inherited variant m.3243A>G mapping in the *MT-TL1* gene encoding the mitochondrial tRNA^Leu(UUR) 2,3^. MELAS can also arise due to additional mitochondrial pathogenic variants with a very low frequency, such as m.1630A>G mapping in the *MT-TV* gene encoding the mitochondrial tRNA^Val 4–6^, m.13514A>G mapping in the *ND5* gene encoding the NADH dehydrogenase 5 subunit of Complex I ^7^, and m.14453G>A mapping in the *ND6* gene encoding the NADH dehydrogenase 6 subunit of Complex I ^8^.

These mitochondrial MELAS variants only affect a subset of the multi-copy mitochondrial genome, a phenomenon called heteroplasmy stemming from a variable ratio of mutant and wild-type mitochondrial DNA (mtDNA) coexisting within cells ^9^. When the population of mutated mtDNAs overwhelms the wild-type mtDNA population, mitochondria become dysfunctional in terms of OXPHOS. Consequently, patients harboring a MELAS variant generally are symptomatic when the load of dysfunctional mitochondria exceeds a certain threshold of heteroplasmy, a threshold that is tissue-specific and varies among patients. This genetic heterogeneity compounded by the unprecedented clinical heterogeneity among MELAS patients has made the genotype-phenotype relationship elusive, and consequently the pathogenic MELAS mechanism ^10^. These hurdles have hampered progress toward curative interventions and reliable biomarkers and therapeutic targets.

Recent advancements in multi-omics analysis have enabled systematic insights into disease processes and correlations of different classes of biomolecules in disease pathogenesis^11–16^. In this study, we designed a global and integrated mass spectrometry (MS)-based strategy for parallel proteomics and metabolomics to provide novel insights into the pathogenic pathways of MELAS. Integrated omics experiments were conducted on dermal fibroblasts derived from a female exhibiting the cardinal symptoms of MELAS. This patient harbored the rare mitochondrial variant m.14453G>A with a 65% heteroplasmy, specifically affecting the mitochondrial-encoded subunit NADH dehydrogenase 6 of the OXPHOS Complex I. This pathogenic variant was not detected in her mother’s fibroblasts, making this *de novo* mitochondrial variant ultra-rare.

The strength of our study for understanding the MELAS mitochondrial pathogenesis via proteomics and metabolomics platforms is two-fold: 1) the ideal *ex vivo* cellular paradigm by pairing dermal fibroblasts from the symptomatic MELAS patient with her non-carrier mother as a negative control; and 2) our comprehensive coverage and mutual verification of identified proteins and metabolites using various modes of LC-MS/MS with a combination of data-dependent (DDA) and data-independent acquisitions (DIA). For the past decades, DDA has been the first and standard strategy in shotgun proteomics, where the most abundant sets of MS1 ions are individually selected and isolated for sequential MS^2^ fragmentation^17–19^. For label-free global proteomics, identification is achieved at MS^2^ level, and quantification is often achieved at MS1 level. However, for complex biological samples, DDA proteomics often faces challenges to identify low abundant and coeluting peptides, as well as missing values from different biological/technical replicates. As an alternative strategy to DDA proteomics, DIA isolates and fragments all MS1 ions within a given *m/z* window regardless of their intensities. Both identification and quantification can be achieved at the MS^2^ level with fewer missing values and often better reproducibility compared to DDA. But DIA generates highly convoluted spectra and requires tailored computational algorithms for data analysis. With the rapid advancements of DIA data analysis, many software platforms became available to analyze DIA data, such as Spectronaut ^20^, Skyline ^21^, DIA-Umpire ^22^, DIA-NN ^23^, and the most recently developed MaxDIA^24^ in the Maxquant platform, to name but a few. In this study, we conducted global DDA proteomics of MELAS fibroblasts vs. controls and verified the findings with DIA analysis. The performance of the newly launched MaxDIA platform was evaluated in comparison to the widely used Spectronaut platform. We report m.14453G>A-specific proteomic and metabolomic fingerprints present in dermal fibroblasts from a symptomatic MELAS patient compared to her asymptomatic mother as a perfect negative control group, revealing pathogenic pathways congruent with the patient’s Complex I deficiency and chronic energy deficit.

## EXPERIMENTAL

### Subject, Skin Biopsy, and Fibroblast Culture

This study was approved by the Institutional Review Board of the George Washington University and Children’s National Medical Center. It was conducted in accordance with the ethical principles of the Declaration of Helsinki of 1975 (revised 1983). Patient skin biopsy was performed only after receiving written informed consent with permission to study the derived dermal fibroblasts. Skin biopsy was performed on a 22-year-old symptomatic female harboring the m.14453G>A variant and her 53-year-old asymptomatic mother as a negative control. Dermal fibroblasts were derived from a 2 mm punch skin biopsy in Dulbecco’s Modified Eagle Medium (DMEM; Gibco) supplemented with 2 mM glutamine, 2.5 mM pyruvate, 0.2 mM uridine, FGF-2 (10 ng/ml) and 20% fetal bovine serum as described in a previous study ^6^. Derived dermal fibroblasts were frozen at passage 2 and never used beyond passage 10.

### Determination of Heteroplasmy

Total DNA was extracted from cultured dermal fibroblasts at passage 3 using the QIAmp DNA mini kit according to the manufacturer’s recommendations (Qiagen). Heteroplasmy was determined using a Long-Range PCR-based Next-generation sequencing approach ^25^. We applied a very stringent detection method by choosing the very stringent cut-off of 1.33% heteroplasmy based on three standard deviation above the mean error, which resulted in a 99.9 % confidence ^26^.

### Lysis of Dermal Fibroblasts and Biomolecule Extraction

Dermal fibroblasts derived from the MELAS patient and her noncarrier mother were grown in four independent biological replicates. Fibroblasts were washed twice with phosphate-buffered saline and immediately pelleted and flash-frozen in liquid nitrogen. Proteins, polar metabolites, and nonpolar metabolites/lipids were enriched from fibroblasts using methanol/chloroform/water extraction ^27,28^. Briefly, 300 μL of ice-cold methanol, chloroform, and MilliQ water were sequentially added to each sample. The biphasic mixture was vigorously vortexed, incubated on ice for 10 min, and clarified by centrifugation at 12,000 rpm for 15 min at 4 °C. The mixture stratified into three layers: the top methanol/water fraction containing polar small molecules, the middle layer of protein precipitate, and the bottom chloroform fraction containing nonpolar metabolites and lipids. Different classes of biomolecules were collected separately in Eppendorf tubes and stored at −80°C.

### Protein Sample Preparation

Protein pellets were dissolved in lysis buffer containing 150 mM NaCl, 50 mM Tris, and 0.1% (w/v) Rapigest (Waters), and sonicated with QSonica Q700 in an ice-cold water bath with alternating cycles of 1min on and 30 s off for a total of 15 min. The protein solution was clarified by centrifugation at 12000 rpm for 10 min at 4 °C. Total protein concentrations of each sample were determined and normalized using DCA colorimetric protein assay (Bio-Rad). Protein reduction and alkylation were performed by adding tris(2-carboxylethyl) phosphine (TCEP) at 5 mM for 30 min at 37 °C in a ThermoMixer, followed by iodoacetamide (IAA) addition at 15 mM in dark for 30 min at 37 °C. Another 5 mM TCEP was added for 10 min to quench excessive IAA. Trypsin/Lys-C mix (Promega) was used for protein digestion (1:30 ratio w/w) for 16 hrs at 37 °C in a ThermoMixer. Digestion reaction was quenched with 10% trifluoroacetic acid until pH < 2, incubated at 37 °C for 45 min, and clarified by centrifugation at 12000 rpm for 10 min at 4 °C to precipitate and remove Rapigest. Peptides were desalted using a Waters C18 96-well extraction plate following the manufacture protocol, dried, and stored at −30 °C until LC-MS/MS analysis.

### Metabolite Sample Preparation

Polar metabolites (methanol/water fraction) and nonpolar metabolites and lipids (chloroform fraction) were dried separately in SpeedVac and reconstituted in 50/50 acetonitrile/water and 50/50 acetonitrile/isopropanol, respectively. Metabolite samples were normalized based on total protein concentrations. Samples were briefly sonicated, clarified by centrifugation at 12000 rpm for 10 min at 4 °C, and stored at −30 °C until LC-MS/MS analysis.

### nanoLC-MS/MS for DDA and DIA Proteomics

A Dionex Ultimate 3000 RSLCnano system coupled with a Thermo Scientific Q-Exactive HFX Orbitrap Mass Spectrometer was used for proteomic analysis. The mobile phase A was 0.1% formic acid (FA) in water, and mobile phase B was 0.1% FA in acetonitrile. Peptides were injected onto an Acclaim PepMap C18 trap column (3 μm, 100 Å, 75 μm × 2 cm) and separated on an Easy-spray PepMap C18 column (2 μm, 100 Å, 75 μm × 75 cm) with a 210 min LC gradient and 55 °C column temperature. The flow rate was 0.2 μL/min. The quadrupole mass filtering was set from *m/z* 380 to 1500 with a resolving power of 120000 (at *m/z* 200 FWHM). For DDA analysis, a top 40 method was conducted with an MS resolving power of 120K and an MS/MS resolving power of 7500. Parent masses were isolated with an *m/z* 1.4 window and fragmented with higher-energy collision dissociation at a normalized collision energy (NCE) of 30%. The maximum injection times (maxIT) were 50 ms for MS and 35 ms for MS/MS. The dynamic exclusion time was 30 s. The automatic gain control (AGC) was 1 × 10^6^ for MS and 2 × 10^5^ for MS/MS. For DIA analysis, the resolving power was 30K, NCE was 30%, and the isolation window was 8Da (staggered). AGC target was 5 × 10^5^, and the maxIT was 20 ms. MS precursor scan was acquired in parallel to the DIA scan with a mass range of *m/z* 380-1500, a resolving power of 60K, maxIT of 30 ms, and AGC of 1 × 10^6^.

### UHPLC-MS/MS for Polar and Nonpolar Metabolomics

A Vanquish Duo UHPLC system coupled with a Thermo Scientific Q-Exactive HFX Orbitrap Mass Spectrometer was used for metabolomic analysis. The same sample was analyzed twice on a reverse phase (RP) and a hydrophilic interaction chromatography (HILIC) column. For RPLC-MS, metabolites were separated using a Luna Omega Polar C18 column (1.6 μm, 100 Å, 100 x 2.1mm) with a 20 min gradient and 30 °C column temperature. RP aqueous buffer was 0.1% FA in water, and RP organic buffer was 0.1% FA in acetonitrile. The flowrate was 0.3 mL/min.

The quadrupole mass filtering was set from *m/z* 70 to 800 with a resolving power of 45000 (at m/z 200 FWHM), operated on positive electrospray ionization (ESI) mode. For HILIC LC-MS, a Kinetex HILIC column (2.6 μm, 100 Å, 150 x 2.1mm) was used with a 37 min gradient and 30 °C column temperature. HILIC aqueous buffer was 95% water, 5% acetonitrile with 10 mM ammonium acetate. HILIC organic buffer was 5% water, 95% acetonitrile with 10 mM ammonium acetate. The flowrate was 0.3 mL/min. MS data acquisition for each replicate was obtained in full MS mode. The quadrupole mass filtering was set from m/z 70 to 1000 with a resolving power of 45000 (at m/z 200 FWHM), operated on both positive and negative ESI mode. The maximum injection times were 50 ms for full MS mode. The automatic gain control (AGC) was 1 × 10^6^ for full MS mode.

Two quality control (QC) samples were created by pooling a small aliquot from each sample for polar and nonpolar fractions. The QC sample was injected in conditions described above but in Top 10 DDA modes to assist with confident metabolite identification. MS resolving power was 45000 and MS^2^ resolving power was 7500, with an isolation window of *m/z* 1.2, NCE of 30%, and dynamic exclusion of 30 s.

Metabolite standard mixture containing 66 metabolites (e.g., amino acids, nucleotides, neurotransmitters, organic acids, and metabolites from metabolic energy pathways) from our in-house metabolomics library was analyzed (20 μM) using the same RPLC-MS and HILIC-MS methods described above. LC-MS/MS analyses of QC samples and metabolite standards were used for metabolite identification based on our previously established metabolite identification flowchart ^29^.

### DDA and DIA Proteomics Data Analysis

The DDA proteomics dataset was analyzed with Thermo Fisher Proteome Discoverer (PD, 2.4.1.15) software with the SequestHT search engine. Swiss-Prot *Homo sapiens* database (reviewed) was used for human protein identification with 1% false discovery rate cut-off. Proteomics contamination database (from Max Planck Institute of Biochemistry) was included as the contamination marker. Known mitochondrial proteins from Uniprot were included as the mitochondrial marker. Trypsin was used as the enzyme with 3 maximum missed cleavages. Methionine oxidation and acetylation of protein N-terminus were included as variable modifications. Cysteine carbamidomethylation was included as a fixed modification. Chromatographic alignment was conducted with a maximum retention time shift of 2 min and a minimum signal-to-noise ratio of 5. Precursor mass tolerance was 25 ppm. The max fold change is set to 500. Data was normalized by total peptide amount with no missing value imputation.

The DIA dataset was analyzed using two software platforms: Spectronaut 15 ^20^ (Biognosys) and recently launched MaxDIA^24^ using MaxQuant (2.0.1.0). The default parameters were used, and the enzyme and modification settings were the same as the DDA data analysis. In Spectronaut, swiss-Prot *Homo sapiens* database (reviewed) was used in library-free directDIA mode. The global imputing function for missing values was turned off. In Maxquant, MaxDIA was used in discovery mode using FASTA file UP000005640_9606 *(H. sapiens)* and *in silico* generated spectral library files as suggested by the MaxDIA software ^24^.

### Metabolomics Data Analysis

Metabolomics dataset was analyzed with Thermo Fisher Compound Discoverer (3.2) software. The maximum retention time shift was 1 min. Mass tolerance was 5 ppm. The minimum peak intensity was 1×10^5^, and the signal-to-noise ratio was 3. DDA acquisitions of QC samples were included for metabolite identification. Four datasets were analyzed separately for polar-RP, polar-HILIC, nonpolar-RP and nonpolar-HILIC data. Positive and negative ESI modes were combined in each dataset. Metabolite identification was conducted based on our previously established flowchart that includes accurate mass matching and MS/MS matching to online metabolite databases as well as spectral matching to the in-house standard library ^29,30^. MzCloud, ChemSpider, HMDB, KEGG, and LIPID MAPS were selected as metabolite libraries in the Compound Discoverer software. The in-house metabolite standard library contained the retention time, MS1, MS/MS, and detected LC mode for 66 common metabolites, such as amino acids, nucleotides, neurotransmitters, organic acids, and metabolites from TCA cycle and glycolysis.

### Statistical analysis and Bioinformatics

Proteomics data were exported in the excel format and further analyzed with R for statistical analysis (t-test) and Spearman’s correlation ^31^. Metabolomics data from 4 different LC modes were combined for further multivariate statistical analysis. Principal component analysis (PCA) and joint pathway analysis were conducted with MetaboAnalyst 5.0 ^32^.

## RESULTS & DISCUSSION

### Advantages of *ex-vivo* Patient-derived Fibroblasts System

We obtained skin biopsies to derive dermal fibroblast from the symptomatic MELAS female and her noncarrier mother as a negative control group. We opted for this *ex vivo* cellular MELAS paradigm because of several advantages ^33,34^: 1) skin biopsy is a non-invasive procedure; 2) the derived fibroblasts can be propagated and stored for future studies to identity personalized biomarkers, genetic markers, and therapeutic targets; 3) heteroplasmy of a specific mitochondrial pathogenic variant in dermal fibroblasts does not decline with age of the patient nor vary between genders, essential to decipher the MELAS pathogenic mechanism by integrating omic-driven phenotypes, heteroplasmic severity, and disease burden ^35^; and 4) fibroblasts provide intact mitochondria in contrast with biofluid samples like blood and urine ^9,36^; 5) fibroblast culture is suitable for high-throughput multi-omics analysis.

### Experimental Workflow of Integrated Proteomics and Metabolomics

The overall workflow is illustrated in **Figure 1**. We simultaneously extracted proteins, polar metabolites, as well as nonpolar metabolites and lipids from the symptomatic patient’s fibroblasts and those of her asymptomatic and noncarrier mother as a control group using the methanol/chloroform/water extraction. Proteomics was conducted with DDA and verified by DIA. We comparatively evaluated the proteomic results from DDA (PD), DirectDIA (Spectronaut) ^20^, and the newly launched MaxDIA^24^ (Maxquant). Metabolomics was performed in both RP and HILIC LC-MS platforms for polar and nonpolar metabolites to achieve a comprehensive metabolome coverage and mutual verification. Additional analyses of pooled samples and metabolite standards ensured the confident identification of metabolites. Quantitative and qualitative information of all identified proteins and metabolites were combined into the same molecular processes and pathways to understand mitochondrial dysfunctions and pathogenesis involved in MELAS.

**Figure 1.**
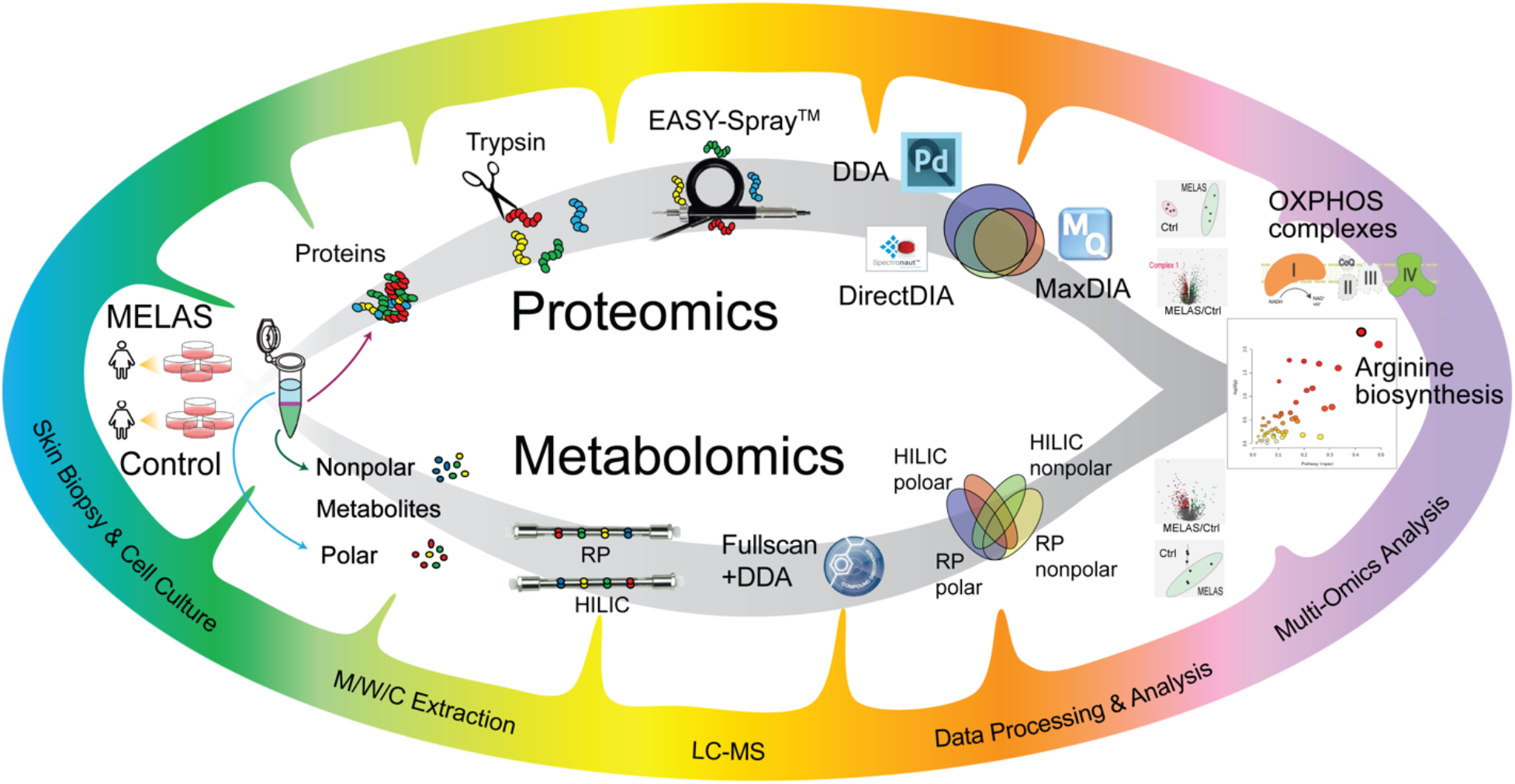
Schematic workflow of integrated MS-based proteomics and metabolomics of MELAS. Proteins, nonpolar metabolites, and polar metabolites were extracted simultaneously from patient and control dermal fibroblast cultures. LC-MS-based proteomics and metabolomics were conducted in parallel and integrated for pathway analysis.

### Comparative Evaluation of DDA and DIA Proteomics

To identify the MELAS proteomic signature specific for the pathogenic m.14453G>A variant, we performed DDA proteomic analysis followed by DIA for verification. After removing contaminants and decoys, a total of 7144 protein groups were identified from human fibroblasts, 65% of which overlapped in at least two datasets from DDA (PD), directDIA (Spectronaut) and MaxDIA (MaxQuant) platforms (**Figure 2A**). Majority of proteins (94%) identified in directDIA overlapped with DDA result, but not with MaxDIA. MaxDIA identified more total proteins but has significantly more missing values compared to directDIA. DDA method generated the most protein IDs (6994) compared to library-free directDIA (4587) and maxDIA (4868). As patient-derived fibroblasts with pathogenic variant were difficult to scale up, we didn’t conduct fractionation or generating DIA spectral library in this study. But for other unlimited sample materials, protein identifications could be further increased by offline LC fractionation or building a comprehensive spectral library ^37,38^. Without using missing value imputation, DirectDIA notably returned minimum missing values (0.02%), whereas DDA and MaxDIA results contained 9.1% and 13.7% missing values, respectively. From these identified protein groups, 5487 from DDA, 4587 from directDIA, and 3141 from MaxDIA were reproducibly quantified in at least 3 replicates in one group (MELAS or control) (**Figure 2B**). Among the commonly quantified 2054 protein groups, 1243 proteins (60.5%) shared a consistent trend (increase or decrease) in MELAS vs. Ctrl groups in all 3 datasets. The complete DDA and DIA proteomics datasets were merged in **Supplemental Table S1**. When we examined the proteins that reach statistical significance (*p-* value <0.05), 174 proteins were significantly altered in all 3 datasets, 170 of which have the same changing trend (**Figure 2C**). Spearman’s correlations of these three data analysis platforms indicated overall consistent protein fold changes **(Figure 2D)**. But discrepancies and outliers do exist in DDA vs. DIA datasets, demonstrating the necessity to verify proteomics results in multiple analytical or data analysis platforms. PCA results of DDA, directDIA, and MaxDIA data all showed complete separation between the MELAS (symptomatic daughter) and the control (asymptomatic mother) groups (**Figure 2E**). While we have extended our data mining using two DIA software, we acknowledge that this is not an exhaustive comparison of DIA methods. Other DIA platforms can also be used here, such as DIA-Umpire^22^, PECAN^39^, Skyline^21^, DIA-NN^23^, etc.

**Figure 2.**
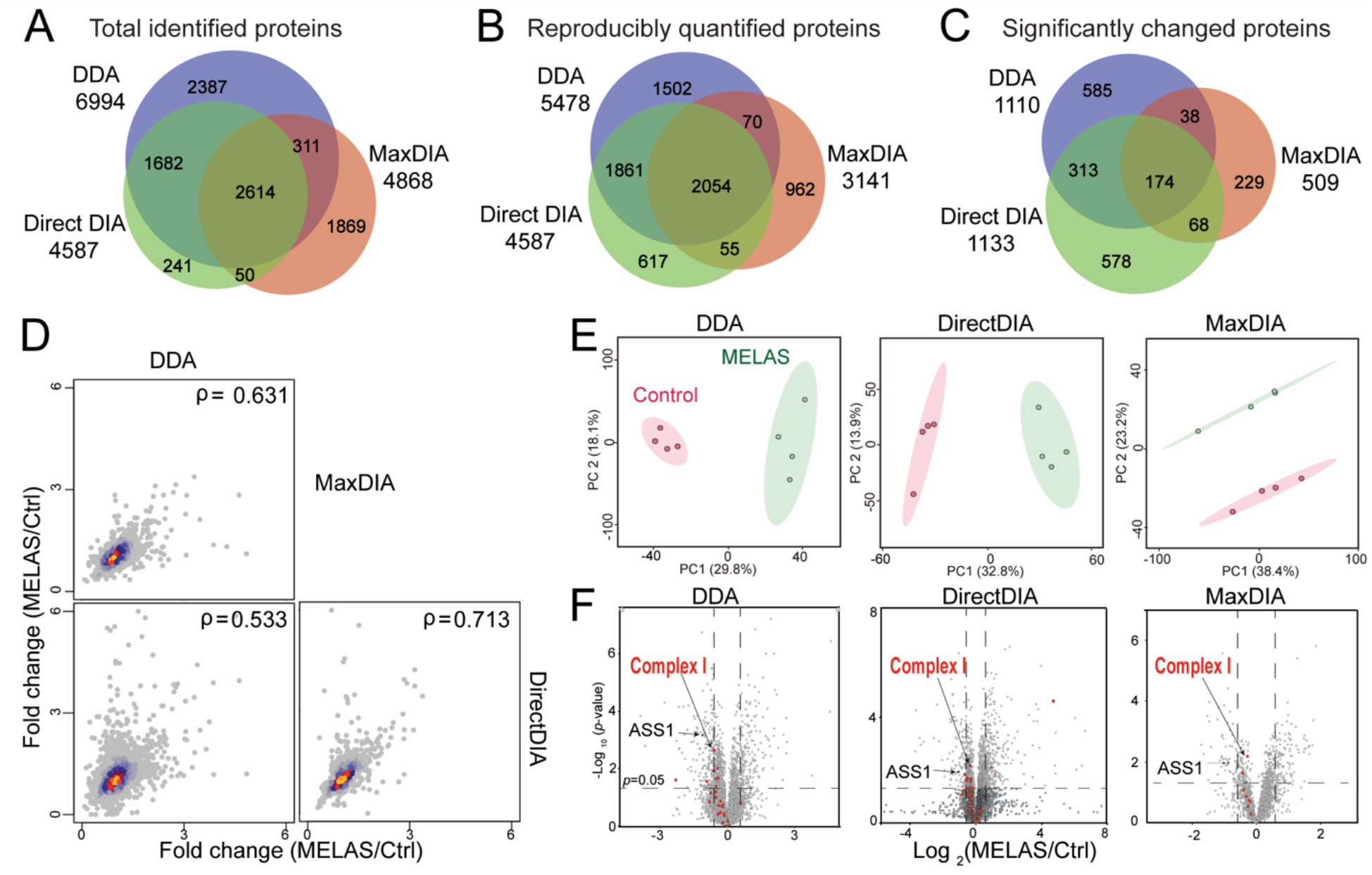
MELAS Proteomic results comparing DDA (Proteome Discoverer), DirectDIA (Spectronaut), and MaxDIA (Maxquant) software platforms. Venn diagrams of three software platforms for total protein identifications (A), reproducibly quantified proteins (B), and significantly changed proteins with *p-*value <0.05 (C). (D) Spearman’s correlation of protein fold changes (MELAS/Ctrl) in DDA, DirectDIA and MaxDIA. (E) Principal component analyses. The ellipses indicate 95% confidence region. (F) Volcano plots of MELAS vs. Ctrl proteomics from three software platforms. Dash lines indicate *p*-value=0.05 and fold change=1.5.

### Global MELAS Proteomics Reveals the Deficits in the OXPHOS Complexes

As the MELAS patient in this study harbors the m.14453G>A variant that specifically affects the OXPHOS Complex I, we highlighted the mitochondrial Complex I subunits identified in our dataset (**Figure 2F**). Indeed, many mitochondrial Complex I subunits showed consistent down-regulation in all three datasets. However, many proteins didn’t reach statistical significance possibly due to the limited sample size (N=4) originated from the difficulty to scale up fibroblast culture from patient with mtDNA mutation. Mitochondrial isolation from cultured fibroblasts via density gradient ultracentrifugation could increase the sensitivity to detect mitochondrial proteins, but such approach cannot be used for patient’s mitochondria with mtDNA mutation, which are very fragile and unable to be enriched with ultracentrifugation. Besides Complex I subunit proteins, we further examined the mitochondrial respiratory chain deficiency in all five OXPHOS complexes (**Figure 3**). The OXPHOS system is composed of a series of distinct respiratory complexes, Complex I to IV, and an ATP synthase, also referred to as Complex V. Mitochondrial ATP synthesis is the result of electron transfer through the first four complexes, with Complex I and Complex II being the two points of entry, and ATP synthesis occurring at Complex V ^40,41^. Electrons are provided by oxidation of the reducing equivalents, NADH and FADH2, originating from the metabolic pathways, glycolysis, the tricarboxylic acid (TCA) cycle, and fatty acid oxidation. Complex I is an L-shaped multi-subunit complex with a hydrophobic membrane arm anchored in the mitochondrial inner membrane that is perpendicularly linked to a hydrophilic peripheral or matrix protruding into the mitochondrial matrix ^42^. The ND6 subunit is required for the proper assembly of Complex I ^43,44^. Among the downregulated expression levels of nuclear-encoded subunits, it is worth highlighting the subunits NDUFS2, NDUFS8, NDUFA9, and NDUFB8, which interact with ND6 during the early steps of the membrane arm assembly ^45^. Equally important is the dysregulation of the two NDUFV1 and NDUFV2 core subunits of the N module, along with the structural subunit NDUFS6, thereby affecting the binding of the reduced agent NADH and consequently accepting electrons by Complex I prior to their transfer to the downstream OXPHOS complexes for ATP synthesis. Subunits of Complex IV were also mostly down-regulated (**Figure 3**). Interestingly, our proteomic results revealed deregulated expression of COX6B1 and NDUFA4, which are required for the last two steps of Complex IV assembly, also known to harbor pathogenic variants linked to mitochondrial diseases with an OXPHOS deficit. Of note is the detected upregulated subunits of Complex V (ATP synthase), a potential compensatory response for the patient’s Complex I deficiency caused by the m.14453G>A variant.

**Figure 3.**
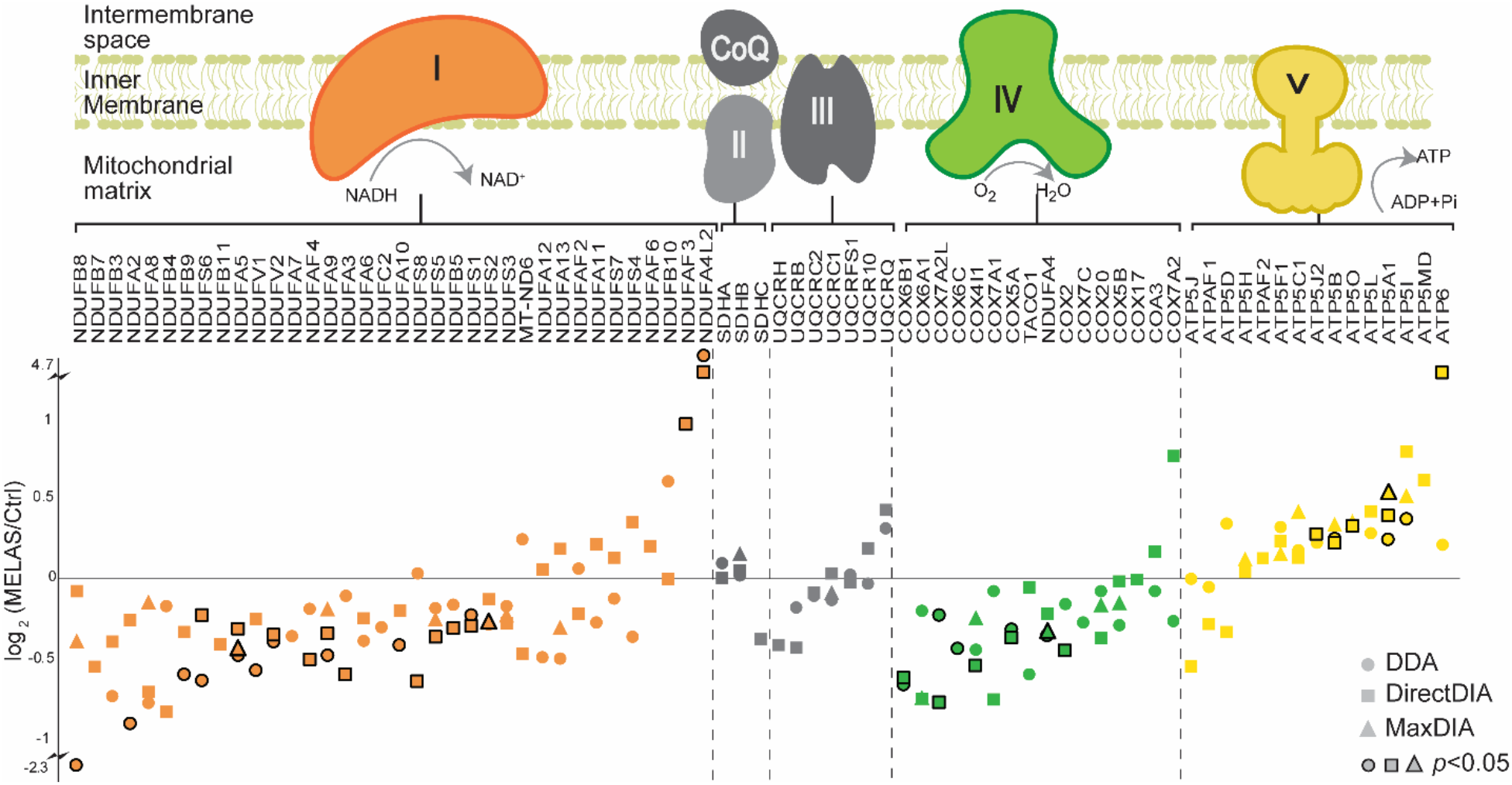
Identified and quantified mitochondrial OXPHOS protein subunits in MELAS vs. Ctrl proteomics. Proteins fold changes from DDA, DirectDIA and MaxDIA were plotted in different shapes. Stroked dots indicate the changes were statistically significant (*p*-value <0.05).

### Comprehensive Metabolomics Revealed Dysregulated Fatty Acid Metabolism in MELAS

Metabolites can reflect the downstream results of endogenous genetic/protein regulations and exogenous influences. MS-based metabolomic technique is particularly useful to understand the molecular processes underlying disease phenotypes and discover potential therapeutic targets for disease diagnosis and management^46–49^. Over 1200 metabolite were identified from a total of ~20K LC-MS peaks from different LC-MS modes (**Supplemental Table S2**). As shown in **Figure 4A**, from the polar metabolite fraction, 661 metabolites were identified in RP mode and 460 metabolites from HILIC mode. From the nonpolar fraction, 262 metabolites were identified in RP and 181 from HILIC. PCA and volcano plots of each LC-MS modes were included in the **Supplemental Figures S1 and S2**. All the ~20K features from different LC modes were then combined to conduct PCA, where complete separation was achieved between the MELAS group (symptomatic carrier daughter) and the control group (asymptomatic and noncarrier mother) (**Figure 4B**). As shown in an example volcano plot in **Figure 4C**, key metabolites related to bioenergetic pathways were altered in MELAS group compared to the control group. The wide range of dysregulated metabolisms may be due to redox imbalance in MELAS patient. Redox imbalance is a direct consequence of the uncoupled electron transport chain ^50^. Deficiency in Complex I is common in MELAS syndrome ^51,52^. Complex I deficit impact not only the electron transport chain, but also the TCA cycle, since Complex I facilitates the conversion of α-ketoglutarate to succinyl-CoA in the TCA cycle. Most notably are the decreased levels of acylcarnitines in the presence of the MELAS m.14453G>A variant, indicative of a defective fatty acid metabolism in the patient-derived fibroblasts. Acylcarnitines modulate the mitochondrial energy metabolism by converting long-chain fatty acids into long-chain acyl-CoAs to overcome the permeability barrier of the inner mitochondrial membrane, thereby allowing their transport into the mitochondrial matrix for fatty acid oxidation and ATP synthesis (**Figure 5B**). This deficiency in acylcarnitine levels further aggravates the chronic energy deficit due to the patient’s Complex I deficiency caused by the m.14453G>A variant.

**Figure 4.**
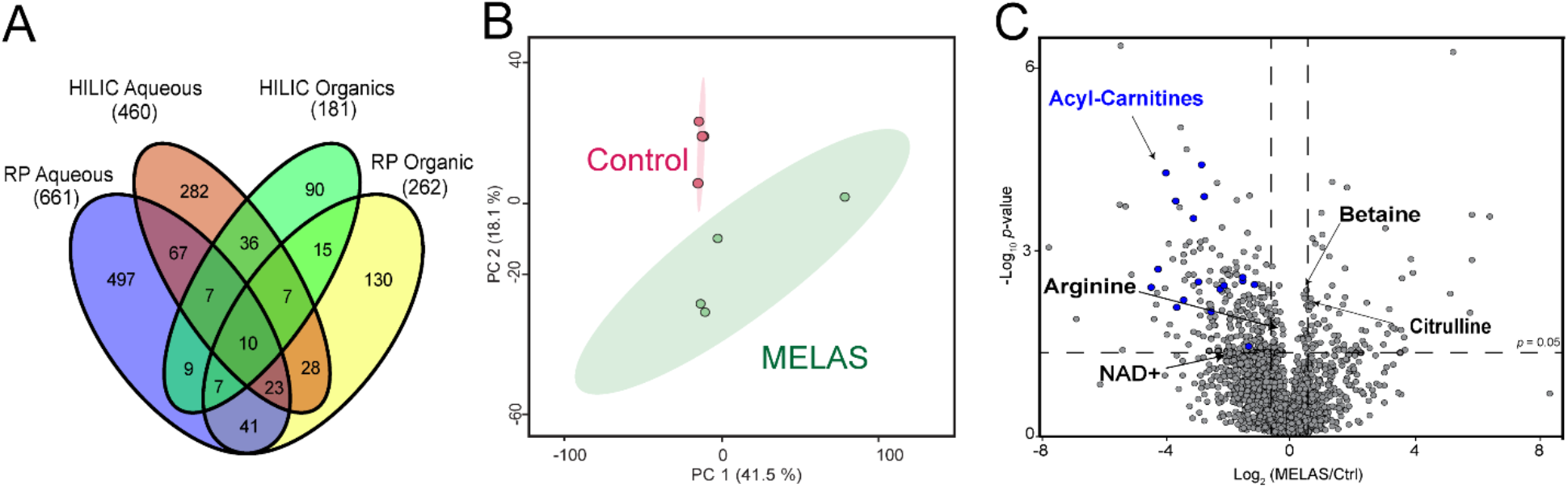
Comprehensive metabolomic fingerprints of MELAS fibroblasts. (A) Venn diagram of identified metabolites from polar/nonpolar fractions using RP or HILIC mode. (B) Principal component analysis from all combined features detected from different LC-MS modes. The ellipses indicate 95% confidence region. (C) Example volcano plot of MELAS vs. control groups from polar metabolites in RP HPLC-MS.

**Figure 5.**
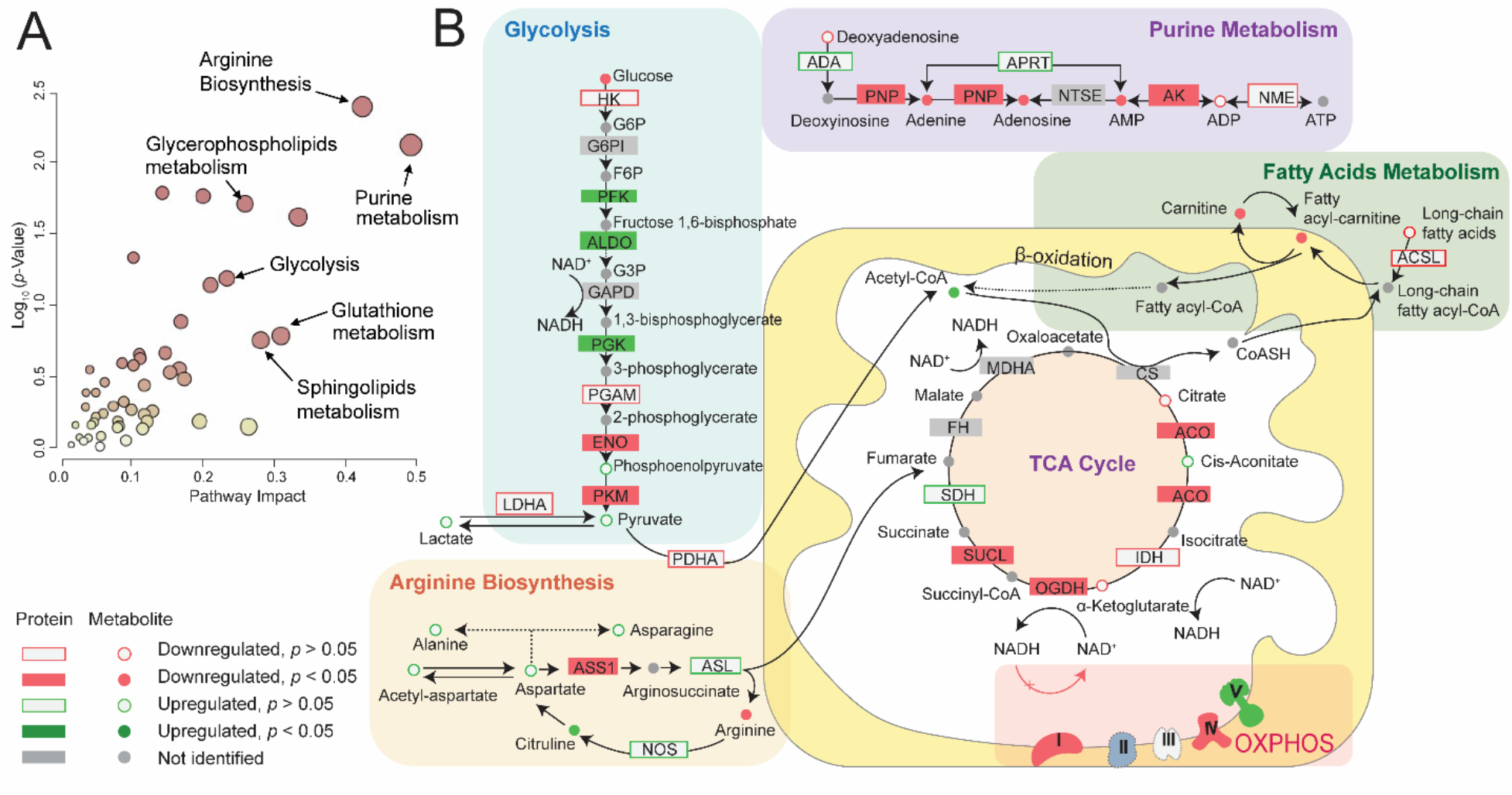
Joint proteomic and metabolomic pathway analysis. (A) Joint pathway analysis results. (B) Highlighted key metabolic pathway segments with both protein and metabolite coverage.

### Multi-Omics Joint Pathway Analysis Highlighted the Deficit in Arginine Biosynthesis in MELAS

To unmask the m14453G>A specific bioenergetic signature, we performed a protein-metabolite joint pathway analysis using the significantly changed proteins and metabolite IDs (**Figure 5A, Supplemental Table S3**). After selecting the significantly altered pathways, all identified metabolites and proteins were included in the pathway nodes despite statistical significance to reveal the overall molecular changes of selected pathways. Among these dysregulated pathways, the arginine biosynthesis pathway has the highest clinical relevance to the MELAS pathophysiology. The key nodal enzyme, arginosuccinate synthase 1 (ASS1), is significantly downregulated, causing the decreased level of arginine (**Figures 2F and 5B**). Decreased plasma levels of arginine has been reported in MELAS patients harboring the most frequent mitochondrial variant m.3243A>G ^1^. Our fibroblast metabolomic analysis also revealed a deficit in arginine level in the context of the m.14453G>A variant, despite the patient’s current treatment of arginine administration (**Figure 4C**). Arginine deficiency may be worsened by the upregulation of nitric oxide synthase (NOS) enzymes, which convert arginine into nitric oxide (NO) and citrulline, a precursor of arginine via the enzymes ASS1 and arginosuccinate lyase (ASL1). The increased levels of citrulline and ASL1 enzyme may represent the compensatory response of cells to regulate arginine biosynthesis but fails to rescue the damage caused by ASS1 deficiency. The prevailing pathogenic mechanism of stroke-like episodes observed in MELAS patients postulates decreased NO availability in the vascular endothelial cells due to endothelial dysfunction causing impaired patency of small cerebral arteries and arterioles ^53,54^. Further aggravating the endothelial NO deficit is the proliferation of dysfunctional mitochondria housing an increased cytochrome c oxidase activity, which promotes binding and sequestration of NO ^55^. Thus, the patient’s arginine biosynthesis fingerprint in our multi-omics study validated the most prevalent clinical manifestations of recurrent stroke-like episodes in this patient. In MELAS patients, stroke-like episodes are nonischemic and the result of impaired blood flow in small intracerebral arteries and arterioles, caused by abnormal proliferation of dysfunctional mitochondria in the vascular endothelial cells and smooth muscle cells in combination of low plasma levels of arginine ^1,56^.

## CONCLUSIONS

In summary, we designed an integrated multi-omics workflow to achieve a comprehensive proteome and metabolome coverage in patient-derived fibroblasts. Our proteomics results provided clues on Complex I deficit, disassembled OXPHOS complexes, and genotype-phenotype correlation of mitochondrial dysfunction caused by the m.14453G>A variant. Impaired mitochondrial energy production is a hallmark of the MELAS pathology, which results in a plethora of phenotypic manifestations targeting several organs. The energy-demanding cells, such as muscle cells and neurons, are particularly vulnerable to such energy deficiency, congruent with two of the cardinal symptoms of MELAS, encephalopathy and myopathy. Our MELAS metabolomics analyses and multi-omics integration revealed dysregulation of fatty acid metabolism, glycerolipid metabolism, sphingolipid signaling pathway, purine metabolism, and glutathione metabolism. Particularly, our multi-omics integration highlighted the dysregulated arginine biosynthesis despite the patient’s daily arginine administration. Decreased arginine level and elevated citrulline level, as well as downregulated ASS1 protein level could serve as potential therapeutic target for treatment. Our findings suggest that the conversion of citrulline into arginine was impeded at the level of ASS1 enzyme. Insufficient arginine level may further aggravate the NO deficiency and therefore the stroke-like episodes in MELAS patients. While this ultra-rare de novo MELAS patient and her healthy mother served as an ideal pair to study the maternally inherited mitochondrial disease, we acknowledge that the biological implications in this study require further validation due to the limited cohort size. Nonetheless, with various highlighted proteins and metabolites, this pilot study set the stage for future MELAS biomarker studies using a larger cohort of MELAS patients.

## Data availability

The proteomics and metabolomics datasets have been deposited in the MassIVE online depository (https://massive.ucsd.edu) with the identifier MSV000088237.

## Author Contributions

L.H., A.C., and H.L. designed the study. H.L., M.U., N.R., A.M., and G.A. conducted the experiments and data analysis. H.L., A.C., and L.H., wrote the manuscript with inputs and revisions from all coauthors. All authors have read and agreed to the published version of the manuscript.

## Acknowledgements

This study is financially supported by the GW Faculty Startup (L.H.), the Department of Defense [W81XWH-20-1-0061] (A.C.) and the NIH National Institute of Child Health and Human Development [1U54HD090257] (A.C.). L.H. acknowledges the ORAU Ralph E. Powe Junior Faculty Enhancement Award. We thank the Vertes lab at GW for the access of the SpeedVac equipment. We thank Ashley M. Frankenfield in the Hao Lab for the assistance with the Spectronaut software.

## Supporting Information

**Figure S1**. Metabolomics principal component analysis plots for each HPLC-MS mode (RP and HILIC, polar and nonpolar metabolites, positive and negative electrospray ionization).

**Figure S2**. Metabolomics volcano plots for each HPLC-MS mode.

**Figure S3**. Example HPLC-MS chromatograms of fibroblast metabolites.

**Table S1**. Merged Protein identification and quantification results from DDA and DIA datasets.

**Table S2**. Merged Metabolite IDs and quantification results from all HPLC-MS modes.

**Table S3**. Joint pathway analysis results.

## References

1 A. W. El-Hattab, A. M. Adesina, J. Jones and F. Scaglia, Mol. Genet. Metab., 2015, 116, 4–12.

2 Y. I. Goto, I. Nonaka and S. Horai, Nature, 1990, 348, 651–653.

3 Y. Kobayashi, M. Y. Momoi, K. Tominaga, T. Momoi, K. Nihei, M. Yanagisawa, Y. Kagawa and S. Ohta, Biochem. Biophys. Res. Commun., 1990, 173, 816–822.

4 M. Bataillard, E. Chatzoglou, L. Rumbach, D. Sternberg, A. Tournade, P. Laforêt, C. Jardel, T. Maisonobe and A. Lombès, Neurology, 2001, 56, 405–407.

5 C. Glatz, K. D’Aco, S. Smith and N. Sondheimer, Mitochondrion, 2011, 11, 615–619.

6 M. Uittenbogaard, H. Wang, V. W. Zhang, L. J. Wong, C. A. Brantner, A. Gropman and A. Chiaramello, Mol. Genet. Metab., 2019, 126, 429–438.

7 P. Corona, C. Antozzi, F. Carrara and L. D’Incerti, Ann. …, 2001, 1, 104–134.

8 K. Ravn, F. Wibrand, F. J. Hansen, N. Horn, T. Rosenberg and M. Schwartz, Eur. J. Hum. Genet., 2001, 9, 805–809.

9 H. Li, M. Uittenbogaard, L. Hao and A. Chiaramello, Metabolites, , DOI:10.3390/metabo11040233.

10 G. S. Gorman, P. F. Chinnery, S. DiMauro, M. Hirano, Y. Koga, R. McFarland, A. Suomalainen, D. R. Thorburn, M. Zeviani and D. M. Turnbull, Nat. Rev. Dis. Prim., 2016, 2, 1–23.

11 M. T. Odenkirk, D. M. Reif and E. S. Baker, Anal. Chem., 2021.

12 C. P. Lapointe, J. A. Stefely, A. Jochem, P. D. Hutchins, G. M. Wilson, N. W. Kwiecien, J. J. Coon, M. Wickens and D. J. Pagliarini, Cell Syst., , DOI:10.1016/j.cels.2017.11.012.

13 D. I. Kim, J. A. Cutler, C. H. Na, S. Reckel, S. Renuse, A. K. Madugundu, R. Tahir, H. L. Goldschmidt, K. L. Reddy, R. L. Huganir, X. Wu, N. E. Zachara, O. Hantschel and A. Pandey, J. Proteome Res., 2018, 17, 759–769.

14 M. T. Odenkirk, K. G. Stratton, L. M. Bramer, B. J. M. Webb-Robertson, K. J. Bloodsworth, M. E. Monroe, K. E. Burnum-Johnson and E. S. Baker, Biomolecules, , DOI:10.3390/biom11010040.

15 L. Hao, J. Johnson, C. B. Lietz, A. Buchberger, D. Frost, W. J. Kao and L. Li, Anal. Chem., 2017, 89, 1138–1146.

16 Y. He, E. H. Rashan, V. Linke, E. Shishkova, A. S. Hebert, A. Jochem, M. S. Westphall, D. J. Pagliarini, K. A. Overmyer and J. J. Coon, Anal. Chem., , DOI:10.1021/acs.analchem.0c04764.

17 R. Aebersold and M. Mann, Nature, 2003, 422, 198–207.

18 C. Fernández-Costa, S. Martínez-Bartolomé, D. B. McClatchy, A. J. Saviola, N. K. Yu and J. R. Yates, J. Proteome Res., 2020, 19, 3153–3161.

19 L. Hao, S. Thomas, T. Greer, C. M. Vezina, S. Bajpai, A. Ashok, A. M. De Marzo, C. J. Bieberich, L. Li and W. A. Ricke, Am. J. Physiol. -Ren. Physiol., 2019, 316, F1236–F1243.

20 R. Bruderer, O. M. Bernhardt, T. Gandhi, S. M. Miladinović, L. Y. Cheng, S. Messner, T. Ehrenberger, V. Zanotelli, Y. Butscheid, C. Escher, O. Vitek, O. Rinner and L. Reiter, Mol. Cell. Proteomics, 2015, 14, 1400–1410.

21 B. MacLean, D. M. Tomazela, N. Shulman, M. Chambers, G. L. Finney, B. Frewen, R. Kern, D. L. Tabb, D. C. Liebler and M. J. MacCoss, Bioinformatics, 2010, 26, 966–968.

22 C. C. Tsou, D. Avtonomov, B. Larsen, M. Tucholska, H. Choi, A. C. Gingras and A. I. Nesvizhskii, Nat. Methods, 2015, 12, 258–264.

23 V. Demichev, C. B. Messner, S. I. Vernardis, K. S. Lilley and M. Ralser, Nat. Methods, 2020, 17, 41–44.

24 P. Sinitcyn, H. Hamzeiy, F. Salinas Soto, D. Itzhak, F. McCarthy, C. Wichmann, M. Steger, U. Ohmayer, U. Distler, S. Kaspar-Schoenefeld, N. Prianichnikov, Ş. Yılmaz, J. D. Rudolph, S. Tenzer, Y. Perez-Riverol, N. Nagaraj, S. J. Humphrey and J. Cox, Nat. Biotechnol., , DOI:10.1038/s41587-021-00968-7.

25 H. Cui, F. Li, D. Chen, G. Wang, C. K. Truong, G. M. Enns, B. Graham, M. Milone, M. L. Landsverk, J. Wang, W. Zhang and L. J. C. Wong, Genet. Med., 2013, 15, 388–394.

26 W. Zhang, H. Cui and L. J. C. Wong, Clin. Chem., 2012, 58, 1322–1331.

27 B. E.G. and D. W.J., Can. J. Biochem. Psychiatry, 1959, 37, 422–422.

28 B. C. Muthubharathi, T. Gowripriya and K. Balamurugan, Mol. Omi., 2021, 17, 210–229.

29 L. Hao, T. Greer, D. Page, Y. Shi, C. M. Vezina, J. A. Macoska, P. C. Marker, D. E. Bjorling, W. Bushman, W. A. Ricke and others, Sci. Rep., 2016, 6, 30869.

30 L. Hao, J. Wang, D. Page, S. Asthana, H. Zetterberg, C. Carlsson, O. C. Okonkwo and L. Li, Sci. Rep., 2018, 8, 1–10.

31 C. Wu, Q. Ba, D. Lu, W. Li, B. Salovska, P. Hou, T. Mueller, G. Rosenberger, E. Gao, Y. Di, H. Zhou, E. F. Fornasiero and Y. Liu, Dev. Cell, 2021, 56, 111–124.e6.

32 Z. Pang, J. Chong, G. Zhou, D. A. De Lima Morais, L. Chang, M. Barrette, C. Gauthier, P. É. Jacques, S. Li and J. Xia, Nucleic Acids Res., 2021, 49, W388–W396.

33 A. Saada, Int. J. Biochem. Cell Biol., 2014, 48, 60–65.

34 M. Uittenbogaard and A. Chiaramello, Mol. Genet. Metab., 2020, 131, 38–52.

35 J. P. Grady, S. J. Pickett, Y. S. Ng, C. L. Alston, E. L. Blakely, S. A. Hardy, C. L. Feeney, A. A. Bright, A. M. Schaefer, G. S. Gorman, R. J. McNally, R. W. Taylor, D. M. Turnbull and R. McFarland, EMBO Mol. Med., 2018, 10, 1–13.

36 S. Thomas, L. Hao, W. A. Ricke and L. Li, PROTEOMICS--Clinical Appl., 2016, 10, 358–370.

37 P. Navarro, J. Kuharev, L. C. Gillet, O. M. Bernhardt, B. MacLean, H. L. Röst, S. A. Tate, C. C. Tsou, L. Reiter, U. Distler, G. Rosenberger, Y. Perez-Riverol, A. I. Nesvizhskii, R. Aebersold and S. Tenzer, Nat. Biotechnol., , DOI:10.1038/nbt.3685.

38 B. C. Searle, L. K. Pino, J. D. Egertson, Y. S. Ting, R. T. Lawrence, B. X. MacLean, J. Villén and M. J. MacCoss, Nat. Commun., , DOI:10.1038/s41467-018-07454-w.

39 Y. S. Ting, J. D. Egertson, J. G. Bollinger, B. C. Searle, S. H. Payne, W. S. Noble and M. J. MacCoss, Nat. Methods, 2017, 14, 903–908.

40 R. Acin-Perez and J. A. Enriquez, Biochim. Biophys. Acta - Bioenerg., 2014, 1837, 444–450.

41 H. Li, A. M. Frankenfield, R. Houston, S. Sekine and L. Hao, J. Am. Soc. Mass Spectrom., , DOI:10.1021/jasms.1c00079.

42 L. A. Sazanov, Nat. Rev. Mol. Cell Biol., 2015, 16, 375–388

43 Y. Bai and G. Attardi, EMBO J., 1998, 17, 4848–4858.

44 E. Perales-Clemente, E. Fernández-Vizarra, R. Acín-Pérez, N. Movilla, M. P. Bayona-Bafaluy, R. Moreno-Loshuertos, A. Pérez-Martos, P. Fernández-Silva and J. A. Enríquez, Mol. Cell. Biol., 2010, 30, 3038–3047.

45 M. Mimaki, X. Wang, M. McKenzie, D. R. Thorburn and M. T. Ryan, Biochim. Biophys. Acta - Bioenerg., 2012, 1817, 851–862.

46 C. H. Johnson, J. Ivanisevic and G. Siuzdak, Nat. Rev. Mol. Cell Biol., 2016.

47 G. J. Patti, O. Yanes and G. Siuzdak, Nat. Rev. Mol. Cell Biol., 2012, 13, 263–269.

48 X. Zang, M. E. Monge and F. M. Fernández, TrAC - Trends Anal. Chem., 2019.

49 M. E. Monge, J. N. Dodds, E. S. Baker, A. S. Edison and F. M. Fernandez, Annu Rev Anal Chem (Palo Alto Calif), 2019, 12, 177–199.

50 H. Li, M. Uittenbogaard, L. Hao and A. Chiaramello, Metabolites, 2021, 11, 233.

51 M. Chol, S. Lebon, P. Bénit, D. Chretien, P. De Lonlay, A. Goldenberg, S. Odent, L. Hertz-Pannier, C. Vincent-Delorme, V. Cormier-Daire, P. Rustin, A. Rötig and A. Munnich, J. Med. Genet., 2003, 40, 188–191.

52 D. Liolitsa, S. Rahman, S. Benton, L. J. Carr and M. G. Hanna, Ann. Neurol., 2003, 53, 128–132.

53 N. Toda and T. Okamura, Pharmacol. Rev., 2003, 55, 271–324.

54 D. J. Green, A. Maiorana, G. O’Driscoll and R. Taylor, J. Physiol., 2004, 561, 1–25.

55 D. M. Sproule and P. Kaufmann, Ann. N. Y. Acad. Sci., 2008, 1142, 133–158.

56 Y. Koga, N. Povalko, J. Nishioka, K. Katayama, N. Kakimoto and T. Matsuishi, Ann. N. Y. Acad. Sci., 2010, 1201, 104–110.

