## Supporting Information for "Integrated Proteomic and Metabolomic Analyses of the Mitochondrial Neurodegenerative Disease MELAS"

Ling Hao

Assistant Professor of Chemistry

### TABLE OF CONTENTS


- **Figure S1**. Metabolomics principal component analysis for each HPLC-MS mode (RP and HILIC, polar and nonpolar metabolites, positive and negative electrospray ionization).
- **Figure S2**. Metabolomics volcano plots for each HPLC-MS mode.
- **Figure S3**. Example HPLC-MS chromatograms of fibroblast metabolites.
- **Table S1**. Merged protein IDs and quantification results from DDA and DIA datasets.
- **Table S2**. Merged Metabolite IDs and quantification results from all HPLC-MS modes.
- **Table S3**. Joint pathway analysis results.

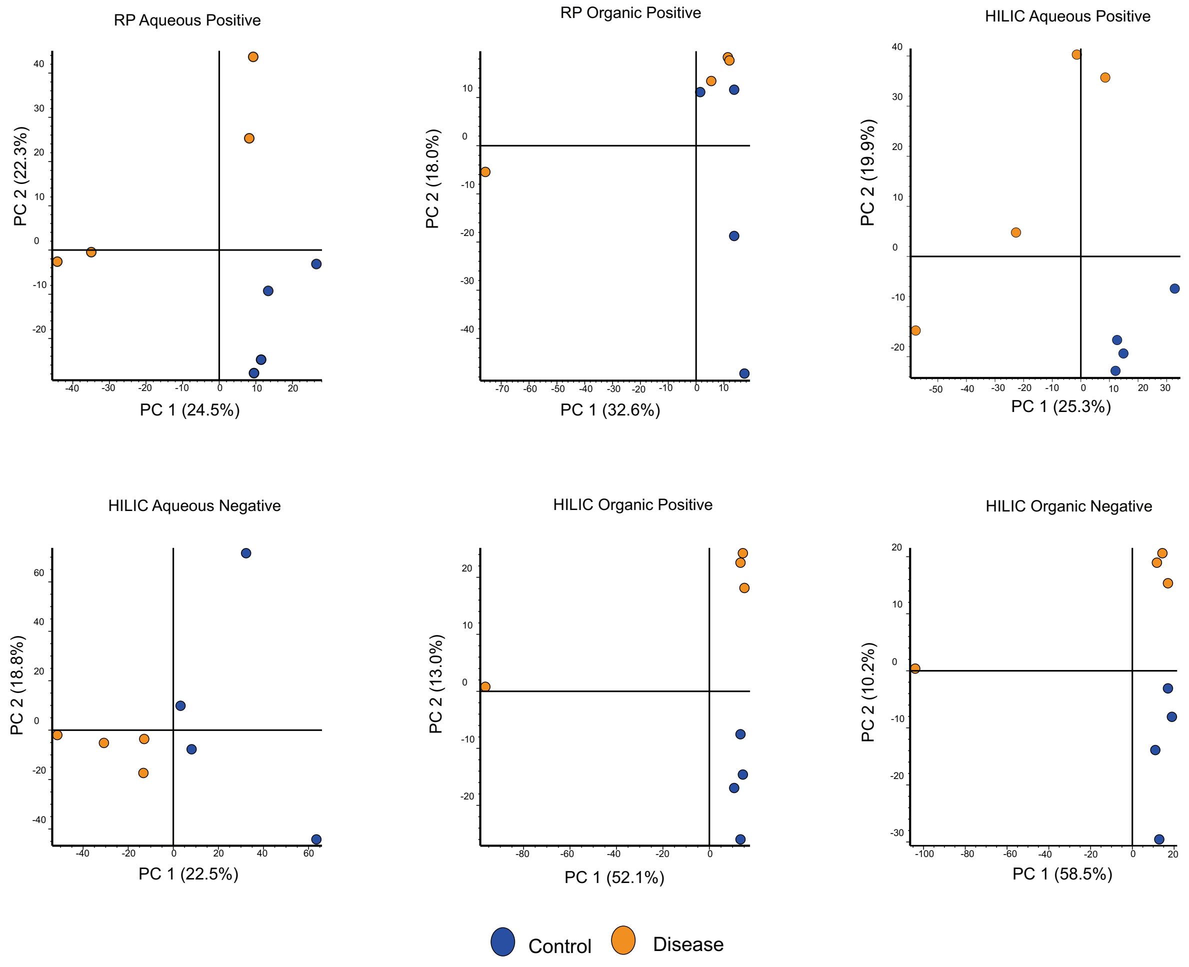


**Figure S1.** Metabolomics principal component analysis for each HPLC-MS mode (RP and HILIC, polar and nonpolar metabolites, positive and negative electrospray ionization).

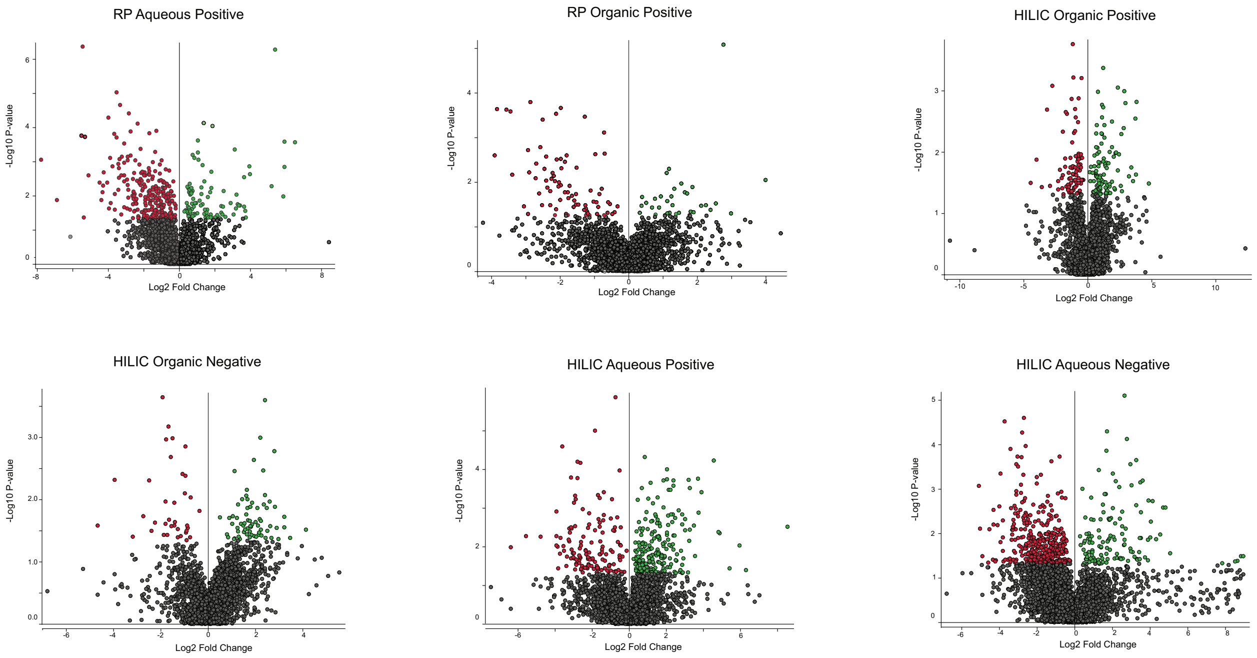


**Figure S2.** Metabolomics volcano plots for each HPLC-MS mode. Aqueous represents the polar fraction from the water/methanol layer, and organic represents the nonpolar fraction from the chloroform layer.

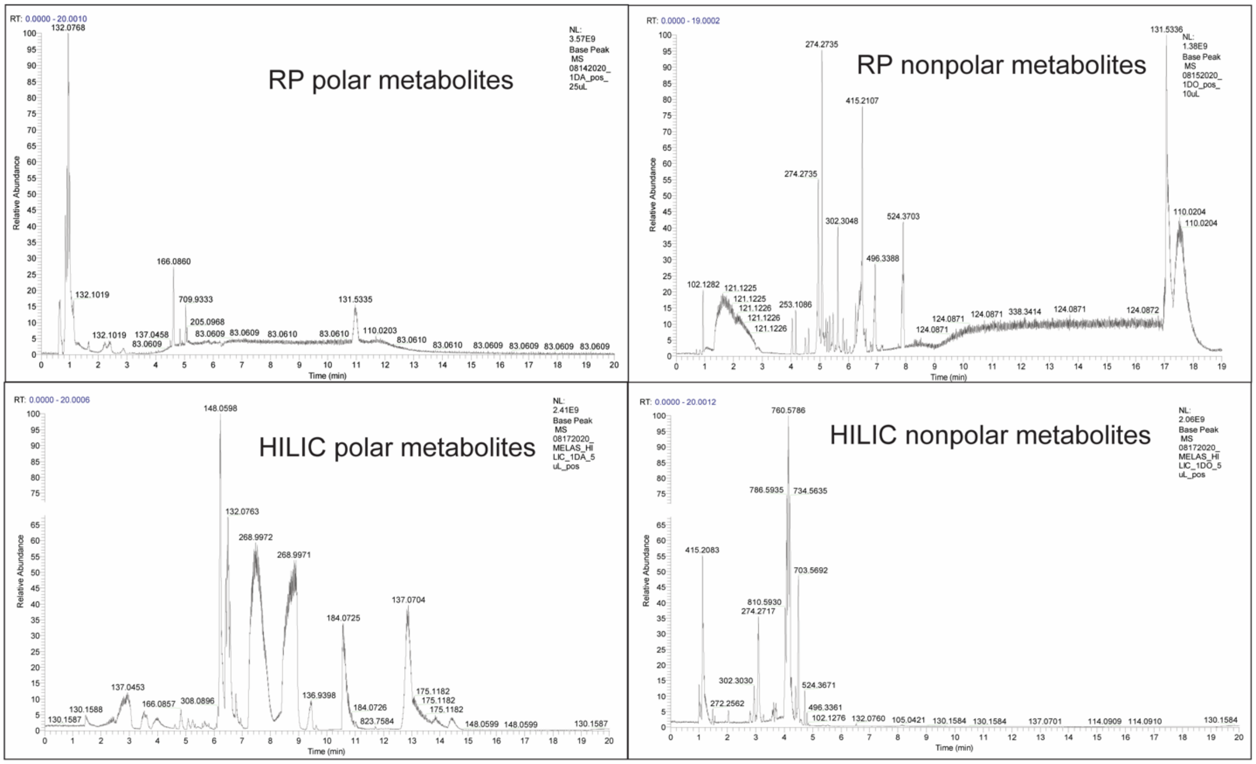


**Figure S3.** Example HPLC-MS chromatograms of fibroblast metabolites.
